# Integration and querying of multimodal single-cell data with PoE-VAE

**DOI:** 10.1101/2022.03.16.484643

**Authors:** Anastasia Litinetskaya, Maiia Schulman, Fabiola Curion, Artur Szalata, Alireza Omidi, Mohammad Lotfollahi, Fabian Theis

**Affiliations:** Institute of Computational Biology, Helmholtz Center Munich, Munich, Germany; School of Life Sciences Weihenstephan, Technical University of Munich, Munich, Germany; Wellcome Sanger Institute, Wellcome Genome Campus, Cambridge, UK; School of Computing, Information and Technology, Technical University of Munich, Munich, Germany; Michael Smith Laboratories, University of British Columbia, Canada; Wellcome–MRC Cambridge Stem Cell Institute, University of Cambridge, Cambridge, UK

**Keywords:** Single-cell, Multimodal integration, Multimodal query-to-reference mapping, Imputation, VAE, Spatial transcriptomic, Metabolomics

## Abstract

Constructing joint representations from multimodal single-cell datasets is crucial for understanding cellular heterogeneity and function. Traditional methods, such as factor analysis and kNN-based approaches, face computational limitations with scalability across large datasets and multiple modalities. In this work, we demonstrate the product-of-experts VAE-based model, which offers a flexible, scalable solution for integrating multimodal data, allowing for the seamless mapping of both unimodal and multimodal queries onto a reference atlas. We evaluate how different strategies for combining modalities in the VAE framework impact query-to-reference mapping across diverse datasets, including CITE-seq and spatial metabolomics. Our benchmarks assess batch effect correction, biological signal preservation, and imputation of missing modalities. We showcase our approach in a mosaic setting, integrating CITE-seq and multiome data to accurately map unimodal and multimodal queries into the joint latent space. We extend this to spatial data by integrating gene expression and metabolomics from paired Visium and MALDI-MSI slides, achieving a high correlation in metabolite predictions from spatial gene expression. Our results demonstrate that this VAE-based framework is scalable, robust, and easily applicable across multiple modalities, providing a powerful tool for data imputation, querying, and biological discovery.

## 1 Introduction

The rapid growth of single-cell technologies has dramatically expanded the availability of multimodal datasets [7]. New methods now enable simultaneous measurement of paired modalities, such as gene expression with ATAC-seq (e.g., 10x multiome [1]), gene expression with protein abundance (e.g., CITE-seq [33]), as well as epigenomic [18] and spatial data [35]. As these technologies advance, the challenge lies in efficiently leveraging these multimodal data in a scalable way.

The rise of multimodal datasets also necessitates foundation models to handle multiple data types simultaneously [9]. This is important for single-cell atlases, which already include millions of cells and will continue to grow [31,23]. These atlases should allow for the mapping of new datasets—both unimodal and multi-modal—onto existing references without requiring re-integration. Furthermore, models must be capable of predicting missing modalities from these atlases, offering a more comprehensive view of cellular states [9].

In this work, we introduce a scalable Product-of-Experts (PoE) Variational Autoencoder (VAE)-based model called Multigrate for multimodal data integration (earlier version presented as a workshop paper at the 2021 ICML Workshop on Computational Biology [24]). Our model integrates datasets with varying numbers of modalities without requiring reimplementation, and it is easily adaptable to new data types for rapid exploratory analysis. It supports the construction of multimodal single-cell atlases, enables both unimodal and multi-modal query-to-reference mapping, and can impute missing modalities. It allows users to map unimodal data onto a multimodal reference and predict missing modalities even when the reference consists entirely of paired multimodal data.

We show that PoE-based integration is more flexible than mixture-of-experts (MoE) models in various query-to-reference settings. Our model outperforms existing approaches on both paired and mosaic integration tasks and successfully predicts spatial metabolomics, measured via MALDI-MSI (matrix-assisted laser desorption/ionization-mass spectrometry imaging) [35], from gene expression data on a query slide. These advancements make our model a robust tool for building and querying multimodal single-cell atlases, offering both scalability and flexibility for advancing biological research.

## 2 Methods

### 2.1 PoE-VAE model

Multigrate is a generative model based on conditional variational autoencoders (cVAEs) [32] and the Product-of-Experts (PoE) approach to combine modalities into a joint representation (Fig. 1A). The architecture consists of three main parts: encoder modules, a PoE module and decoder modules. The input to the model is multimodal single-cell data and the batch covariate (which can indicate, e.g., different datasets, different technical batches or different technologies). The input data is first fed into the encoders separately per modality, which output parameters of unimodal marginal distributions. Then, a product-of-experts module calculates the joint distribution parameters from the marginal distributions’ parameters. We sample from the joint distribution in the latent space and then feed the latent embeddings to the decoders (concatenated with batch covariates). Decoders learn the parameters of the distributions assumed for the input data (e.g., negative binomial for RNA-seq counts).

**Fig. 1.**
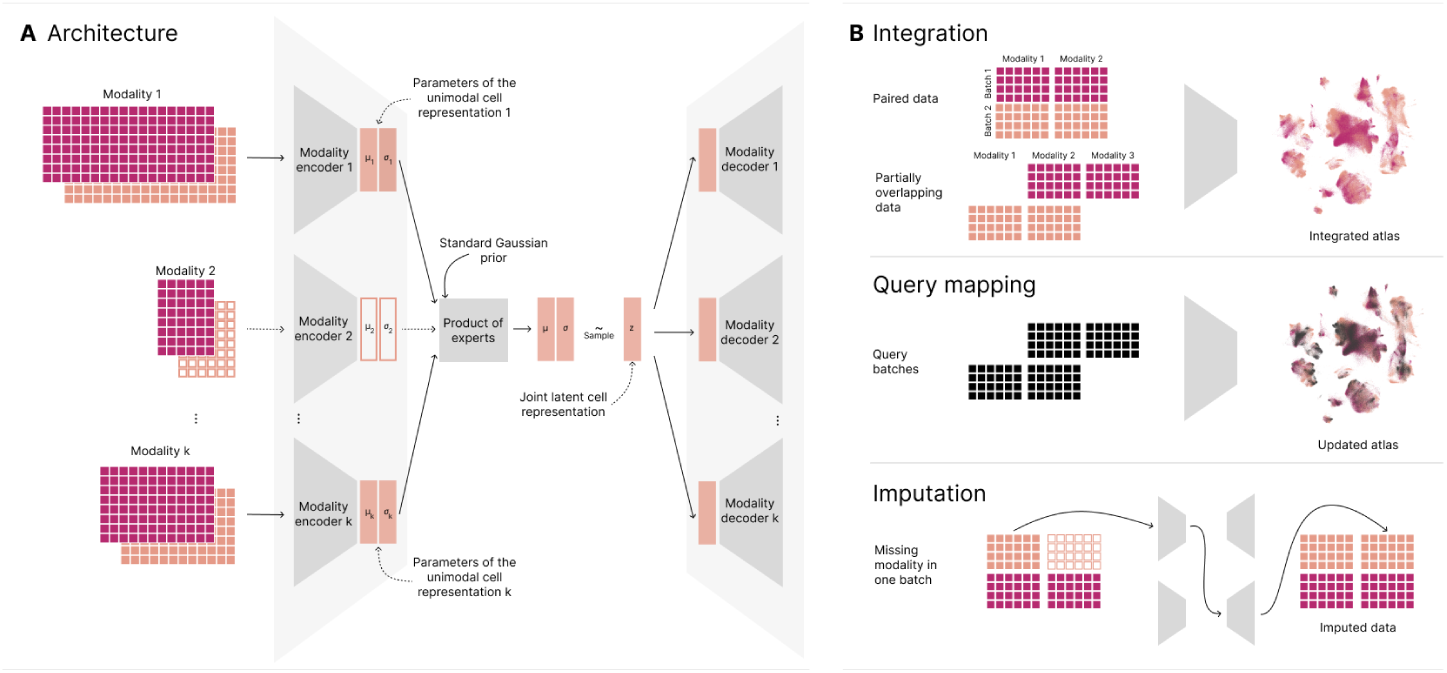
Multigrate enables multimodal integration, query mapping and imputation of missing modalities. **(A)** The Multigrate model accepts paired or partially overlapping single-cell multimodal data as input and consists of pairs of encoders and decoders, where each pair corresponds to a modality. Each encoder outputs a unimodal representation for each cell, and the joint cell representation is calculated from the unimodal representations with Product-of-Experts (PoE). The joint latent representations are then fed into the decoders to reconstruct the input data. **(B)** The key applications for Multigrate are the integration of paired and partially overlapping data into reference atlases (top), mapping of query batches (middle), and prediction of missing modalities (bottom).

Each encoder layer consists of a linear layer with dropout, layer normalization and a non-linearity, which can be chosen by the user (with leaky ReLU as default). The output of the encoders are the parameters of *p*(*z*|*x*_1_)*, …, p*(*z*|*x_m_*), respectively, which are assumed to be normal, and where *x*_1_*, …, x_m_* are unimodal input matrices from *m* modalities. Hence, the output is the mean and variance of each distribution: (*µ*_1_*, σ*_1_)*, …,* (*µ_m_, σ_m_*), where *µ*_1_*, σ*_1_*, …, µ_m_, σ_m_* ∈ ℝ*^n^*^×^*^h^*, *n* is the number of cells in the mini-batch and *h* is the number of latent dimensions. Each parameter is learned independently for each latent dimension. The decoders mirror the encoders’ architecture and consist of blocks of a linear layer with dropout, layer normalization and non-linearity.

We employ the product-of-experts (PoE) [16,22] technique to determine the parameters of the joint distribution *p*(*z*|*x*_1_*, …, x_m_*) from *p*(*z*|*x*_1_)*, …, p*(*z*|*x_m_*) for cell *j* and latent dimension *p* as 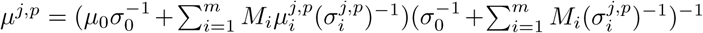 and 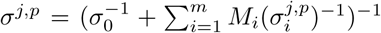, where *µ*_0_ and *σ*_0_ are the parameters of the prior *N* (*µ*_0_*, σ*_0_), which in our case is standard normal, so *µ*_0_ = 0 and *σ*_0_ = 1, and *M_i_* is 1 if modality *i* is present in this particular batch and 0 otherwise. We obtained the closed form above because we assumed all the distributions to be normal [22].

Another approach to calculating the joint representation is with mixture-of-experts (MoE) [29,12]. As we work in the VAEs framework and are interested in the parameters of the joint distribution, we cannot use the more standard formulation of MoE that assumes that 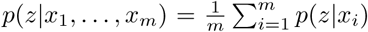, as the resulting distribution is no longer normal, which violates the assumptions on the ELBO loss of the standard VAE. Instead, we assume that the joint latent random variable is an average of unimodal latent variables 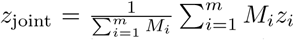, where *z_i_* are drawn independently from *p*(*z*|*x_i_*). This formulation ensures that the joint distribution is again normal with the parameters 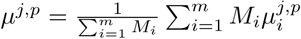 and 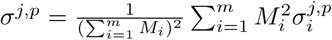. Then, we sample the joint representation *z*_joint_ ∼ *p*(*z*|*x*_1_*, …, x_m_*) independently for each latent dimension using the reparametrization trick [19].

Next, we briefly discuss the maximum mean discrepancy (MMD) loss [14,26]. We employ MMD loss for two purposes: to ensure that the batches are well integrated, i.e., that joint distributions are similar between batches, and that the unimodal representations follow similar distributions. We are interested in the latter if we want to map unimodal queries onto the multimodal reference. MMD loss measures the distance between two distributions *P* and *Q* [14] as MMD(*P, Q*) = 𝔼*_a,a_′*_∼_*_P_* [*K*(*a, a*^′^)] + 𝔼*_b,b_′*_∼_*_Q_*[*K*(*b, b*^′^)] − 2𝔼*_a_*_∼_*_P,b_*_∼_*_Q_*[*K*(*a, b*)], where *a, a*^′^ and *b, b*^′^ are samples drawn from the distributions *P* and *Q*, respectively, and *K* is a kernel function. In the implementation, we use multi-scale radial basis kernels [26] defined as 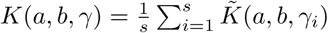, where 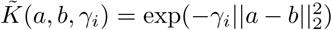 is a Gaussian kernel and *s, γ* = (*γ*_1_*, …, γ_s_*) are hyperparameters.

In our case, the MMD loss is calculated either as the sum over all pairs of batch distributions or as the sum over all pairs of unimodal distributions we want to align. In the first case, MMD loss is calculated between pairs of joint representations 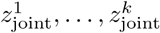 coming from different batches *c*_1_*, …, c_k_* as 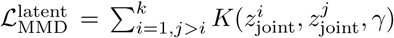. In the second case, we calculate the loss between unimodal marginal representations *z_i_* ∼ *p*(*z*|*x_i_*) and *z_j_* ∼ *p*(*z*|*x_j_*) for all *i, j* ∈ {1*, …, m*}*, i* ≠ *j* as 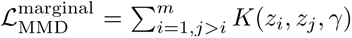. The final MMD loss is calculated as 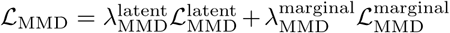, where 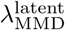 and 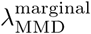 are hyperparameters.

### 2.2 Paired integration

We benchmarked five methods for paired integration (Multigrate, totalVI [13], multiVI [4], MOFA+ [2] and Seurat v4 [15]) on two CITE-seq datasets (NeurIPS 2021 CITE-seq [27], Hao et al. [15]) and two multiome datasets (NeurIPS 2021 multiome [27], 10x multiome [1]). All methods perform multimodal integration of paired data but employ different approaches. MOFA+ is a linear factor model that decomposes the input data into two low-rank matrices, one representing latent factors (i.e., cell embeddings) and the other representing factor effects. WNN is a graph-based method that outputs a nearest-neighbor graph learned from both modalities. totalVI/multiVI are deep-learning VAE-based methods that model and then fit protein-/chromatin-specific distributions. The output of both models is a latent representation in low-dimensional space. We performed hyperparameter optimization for Multigrate and then set Multigrate’s default parameters for the integration task based on the best-performing values across all datasets (Suppl. Methods). Other methods were run with their default parameters. We utilize metrics from [28] (scIB metrics) to evaluate the quality of the integration for all methods. These metrics assess how well the technical noise was corrected and how well the biological signal was preserved in the integrated data. Note that for Seurat v4, we obtained the supervised PCA (sPCA) [6] embeddings from the gene expression and the weighted-nearest neighbor graph to calculate the embedding-based metrics.

### 2.3 Query-to-reference mapping of unimodal data onto the multimodal reference and imputation of missing modalities

We benchmarked how well PoE and MoE approaches impute missing modalities in two different settings (Suppl. Fig. 3A). In the first setting (we call it “query-in-train”), the train set consists of paired data as well as unimodal (usually RNA-seq) data. Then, the model is trained to integrate multimodal and unimodal data in the latent space. To obtain the imputed measurements, the unimodal data is fed through the model to reconstruct the missing modality. The second setting (“no-query-in-train”) does not require unimodal data to be present in the reference. Instead, the model learns to align unimodal representations of the paired data during training. Then, the query data is mapped onto the reference with transfer learning, where only the new weights corresponding to the query data are fine-tuned [25]. After fine-tuning, the query data is passed through the network to reconstruct missing modalities as above. We use the NeurIPS 2021 CITE-seq dataset to predict the protein abundance for the query site and Vicari et al. [35] dataset to predict MALDI-MSI measurements for the query slide. To find the optimal hyperparameters for all the models, we performed a random grid search (Suppl. Methods) and selected the best-performing model based on integration (assessed with scIB metrics) and imputation (Pearson and Spearman correlation).

### 2.4 Mosaic integration and query-to-reference mapping

Mosaic integration refers to the task of integrating datasets that have more than two modalities in total and where some modalities are missing for some of the datasets [3]. For instance, the combined (CITE-seq and multiome) NeurIPS dataset is a mosaic dataset: protein measurements are only available for the CITE-seq batches, ATAC measurements–only for multiome batches, but gene expression is available for all (Fig. 2A).

**Fig. 2.**
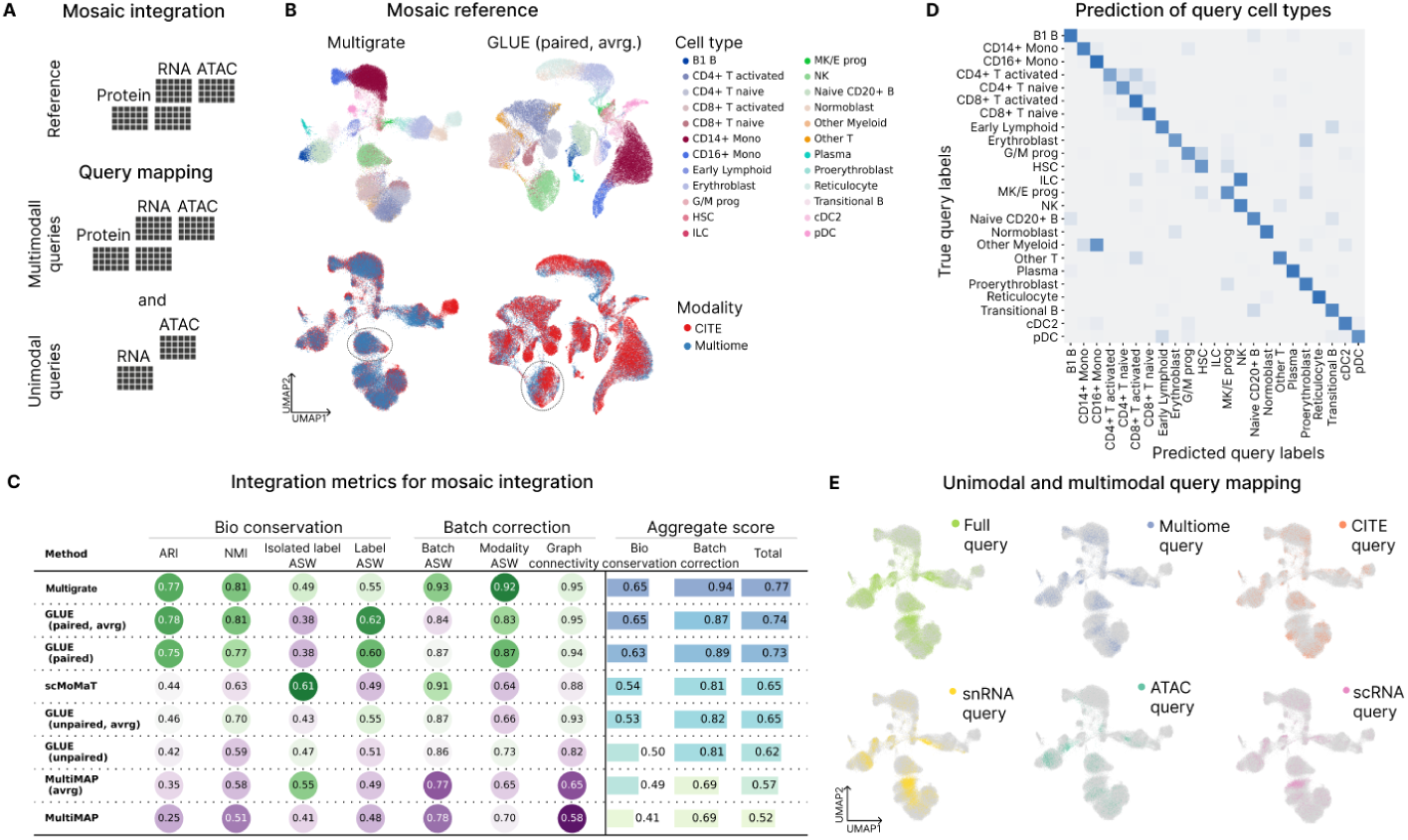
Multigrate enables mosaic integration and query mapping. **(A)** The reference consists of a paired RNA-ATAC and a paired RNA-protein data. The queries are multimodal RNA-ATAC and RNA-protein and unimodal RNA and ATAC data. **(B)** UMAPs of the reference latent space obtained from the two top-performing models (Multigrate on the left and paired GLUE, averaged representation on the right), colored by cell type and modality. **(C)** A table with integration metrics with all the benchmarked methods, showing individual metric scores, averaged bio-conservation and batch-correction scores, and overall scores. **(D)** A confusion matrix between true and predicted (with a random forest model) cell types for the full query mapped with Multigrate. **(E)** UMAPs of different queries mapped onto the trimodal reference with Multigrate.

We benchmarked Multigrate against GLUE [10], multiMAP [17] and scMoMat [39] on this integration task. We subset the combined NeurIPS dataset to Site1 and Site2 and integrated the two batches. We ran GLUE using paired and unpaired models. GLUE offers two different models to train, one that considers the pairedness of the data points and one that does not (Methods); we included both models in our benchmark. Multigrate and scMoMaT output one embedding per cell, while other methods output an embedding per cell per modality. To be able to fairly compare the methods, we additionally computed a “joint” representation for each cell as the average of the modality representations for both of the GLUE models and MultiMAP (denoted “avrg.”).

We mapped unimodal queries, namely scRNA-seq, snRNA-seq and scATAC-seq, and multimodal queries, namely CITE-seq and multiome, on top of the built CITE-multiome reference with Multigrate. We performed a hyperparameter search to report the best-performing model (Suppl. Methods).

### 2.5 Datasets

#### NeurIPS 2021

NeurIPS datasets consist of bone marrow mononuclear cells from healthy donors. The CITE-seq (paired scRNA-seq and surface protein counts) dataset contains 90,261 cells from four sites and 12 batches. The multiome (paired snRNA-seq and scATAC-seq) has 69249 cells from four sites and 13 batches. Both datasets were annotated by the authors and assigned in 30 and 22 cell types, respectively. ‘Samplename’ was used as the batch covariate in the paired and mosaic integration experiments. ‘Site’ was used as the batch covariate in the paired query-to-reference experiments as one of the sites was used as the query.

#### 10x multiome

The data contains 10,000 healthy peripheral blood mononuclear cells (PBMCs) from a multiome experiment. The data does not contain any batches, and the cells are assigned to 11 cell types.

#### Hao et al

The CITE-seq data contains 149,926 PBMCs from eight donors enrolled in an HIV vaccine trial, split into two batches. We used the second-level cell type annotations provided by the authors to calculate the scIB metrics. All 228 proteins were used in the analyses.

#### Vicari et al

This dataset contains slides from coronal sections of a mouse brain. Each slide was processed with MALSI-MSI measuring metabolites (mass/charge) and then processed with Visium to measure spatial transcriptomics (STR). Lipids, neurotransmitters and metabolites were captured using different MALDI matrices, allowing a more diverse description of metabolic processes in the mouse brain but limiting our ability to train one model on all slides due to different feature sets for different slides. We subset the data to 2 slides where the norharmane MALDI matrix was used to measure lipids, namely ‘V11L12-038_A1’ (used as the train set) and ‘V11L12-038_B1’ (used as the query set). We used ‘slide’ as the batch covariate to calculate the quality of the query mapping with scIB metrics as one of the slides was used as the query. We used Leiden clusters [34] calculated on gene expression as ‘label’ for scIB metrics.

#### Data preprocessing

For the paired integration experiments, we subset the gene expression datasets with `scanpy` to the top 4000 highly variable genes, taking the batch covariate into account for datasets with batch effects. If the methods required normalized counts as input, we followed standard `scanpy` workflow and applied log-normalization to the raw counts. Gene expression counts from the Visium experiment were preprocessed the same. We normalized the MALDI-MSI measurements similarly to the gene expression counts and subset the feature space to the top 500 highly variable lipids. We hypothesize that new preprocessing methods will be developed for MALDI-MSI as this data does not contain raw discrete counts but intensities. Protein counts were central-log-ratio normalized. We selected the top 40,000 highly variable peaks for ATAC data with `episcanpy` [11]. To normalize ATAC measurements, we used log-normalization following the `episcanpy` and `muon` tutorials. In the mosaic experiments, we performed the same preprocessing but subsetting to 20,000 highly variable peaks. In the paired query-to-reference mapping experiments, we used 2,000 highly variable genes.

Even though Visium and MALDI-MSI experiments were run on the same slide, we had to align the output of both technologies to obtain the matching between spots to use this data as paired data. We performed the alignment with Moscot [20] using the spatial coordinates as input. After alignment, we obtained 2681 matched spots for slide ‘V11L12-038_A1’ and 2937 matched spots for slide ‘V11L12-038_B1’.

## 3 Results

### 3.1 PoE-VAE model for multimodal single-cell data

Multigrate is a deep-learning model based on conditional variational autoencoders (cVAEs) and allows the integration of multimodal single-cell data, query mapping on new datasets (unimodal and multimodal) and imputation of missing modalities (Fig. 1A,B).

The autoencoder module is implemented as encoder-decoder pairs, where each pair corresponds to a modality present in the data (Fig. 1A). The encoders output the parameters of the corresponding unimodal marginal distribution, and the joint distribution in the latent space is modeled using the Product-of-Experts (PoE) [16,22]. The PoE distribution preserves unique and shared information from the unimodal marginal distributions [22]. The PoE approach also allows Multigrate to integrate paired as well as partially overlapping data (i.e., where the measurements are missing for one or more modalities in part of the data). Additionally, categorical and continuous sample covariates, e.g., technical batch, can be incorporated into the model to obtain the latent representation disentangled from the specified covariates (Methods).

Equipped with scArches transfer-learning approach [25], Multigrate enables query mapping of new datasets onto the multimodal references. The model can accommodate both references trained only on multimodal data or both unimodal and multimodal data, making Multigrate an adaptable tool for multi-modal single-cell analysis.

### 3.2 Multigrate integrates paired multimodal single-cell data

We benchmarked Multigrate’s performance on paired integration against three state-of-the-art methods on two CITE-seq datasets (NeurIPS 2021 CITE-seq [27], Hao et al. [15]) and two paired RNA-ATAC datasets (NeurIPS 2021 multiome [27], 10x public multiome [1]). We compared Multigrate to MOFA+ [2], Seurat v4 WNN [15] on all four datasets, totalVI [13] on CITE-seq datasets and multiVI [4] on the multiome datasets.

To quantitatively evaluate the results, we calculated a subset of the scIB metrics [28] suitable for multimodal integration (Suppl. Methods). The metrics address both the conservation of biological signal and batch effect removal. Overall, Multigrate achieved the highest total score on both paired RNA-ATAC datasets while scoring first and second on the CITE-seq datasets (Suppl. Fig. 3B). TotalVI and Seurat WNN obtained high scores on all datasets, while the score for MultiVI was dataset-dependent (Suppl. Fig. 1). MOFA+ failed to remove batch effects present in the original data, resulting in a low batch correction score (Suppl. Fig. 1, Suppl. Fig. 2).

### 3.3 Multigrate maps unimodal data onto the multimodal reference and imputes missing modalities

We benchmarked the performance of PoE-VAE model against the MoE-VAE model on two datasets: NeurIPS 2021 CITE-seq data [27] and Vicari et al. MALDI-MSI data [35]. We compared how each model performs in each of the query-to-reference mapping scenarios, i.e., ‘query-in-train’ and ‘no-query-in-train’ (see Methods and Suppl. Fig. 3A) and evaluated the mapping with scIB metrics. We also assessed the missing modality prediction with Pearson and Spearman correlation coefficients. Both models achieved very similar correlation on the MALDI-MSI dataset, but the PoE model performed better for the CITE-seq dataset and scored higher on scIB metrics in three out of 4 settings (Suppl. Fig. 3C,D). We, therefore, proceed with PoE as the default for Multigrate and suggest PoE to be the default approach for learning joint representations from multimodal single-cell data.

### 3.4 Multigrate allows mosaic integration and unimodal and multimodal query-to-reference mapping

To demonstrate Multigrate’s ability to perform mosaic integration, we integrated Sites 1 and 2 from the NeurIPS 2021 CITE and Neurips 2021 multiome datasets [27]. We compared Multigrate with GLUE [10], MultiMAP [17] and scMoMaT [39] on this task. We calculated the scIB score on the latent space after performing minimal cell type harmonization between the datasets. We included two Adjusted Silhouette Width (ASW) scores for batch correction: Batch ASW and Modality ASW. This dual-level evaluation of batch and modality mixing allows us to measure the removal of technical biases at a finer scale of individual batches and a coarser scale of modalities simultaneously, aligning with the approach outlined in [21]. For the methods that output one representation per cell per modality, we calculated the metrics once on the original output and once on the averaged representations (denoted “avrg.” in Fig. 2C).

Multigrate scored first, and GLUE (paired model, avrg.) scored second on this task. UMAPs of the learned representations are relatively similar for these two methods (Fig. 2B). Multigrate obtained a slightly higher Modality ASW score than GLUE, which is caused, for instance, by better integrated Natural Killer (NK) cells across modalities (Fig. 2B,C). Overall, we noted that the models that do take into account the information about which cells are paired (Multigrate, GLUE paired) performed better than the methods that do not (Fig. 2C, Suppl. Fig. 4).

When Multigrate’s reference model is trained on multimodal data, our model enables unimodal and multimodal query mapping, where unimodal query modalities can be any of the individual modalities from the multimodal reference. After we build the atlas described above, we map unimodal (i.e., scRNA-seq, snRNA-seq and scATAC-seq) and multimodal (CITE-seq and multiome) queries onto the reference. We calculated scIB metrics using reference and query as two batches to assess the mapping quality. Multigrate successfully mapped all the queries, obtaining very similar scIB scores for all of them (Fig. 2E, Suppl. Fig. 5C). Multimodal queries obtained the highest Batch ASW scores, indicating that the mapping works best for the data modalities present in the reference. We also trained a random forest classifier to transfer the cell types from the reference to the queries and calculated the prediction accuracy. Label transfer worked best for CITE-seq and scRNA data while mapping scATAC-seq seems to be most challenging (Suppl. Fig. 5C,D).

To assess the robustness of our model, we performed several experiments benchmarking the model’s sensitivity towards the number of shared features, the strength of the integration parameter, the size of the reference and the type of the MMD loss (Suppl. Methods and Suppl. Fig. 6). When the number of shared genes is more than 1,000, Multigrate can successfully build the reference, but the quality of query mapping increases with the number of shared features. We also observed that the quality of the query mapping slightly increases with bigger references and that the ‘marginal’ formulation of the MMD loss (see Methods) performed best for unimodal queries.

### 3.5 Multigrate predicts the distribution of spatial metabolites for a query slide

MALDI-MSI is a recent technology that allows users to obtain metabolomic measurements with MSI, H&E (hematoxylin and eosin) staining and STR for the same slide [35]. If we can reliably predict metabolomic intensities from STR, not only can we save time and cost for such experiments but also obtain metabolomic measurements for dozens of existing STR datasets [37,38].

First, we spatially aligned STR and MALDI-MSI slide (Fig. 3A). Then, we matched the aligned spots and fed them as paired data to Multigrate. Our model successfully integrated the paired train slide and the STR query slide, achieving 0.943 Pearson and 0.866 Spearman correlation on the top 500 highly variable lipids. Multigrate learns a joint latent space for spots that captures structure from both RNA and MSI modalities (Fig. 3B). Multigrate can successfully predict different spatial patterns of distribution of metabolites (Fig. 3C).

**Fig. 3.**
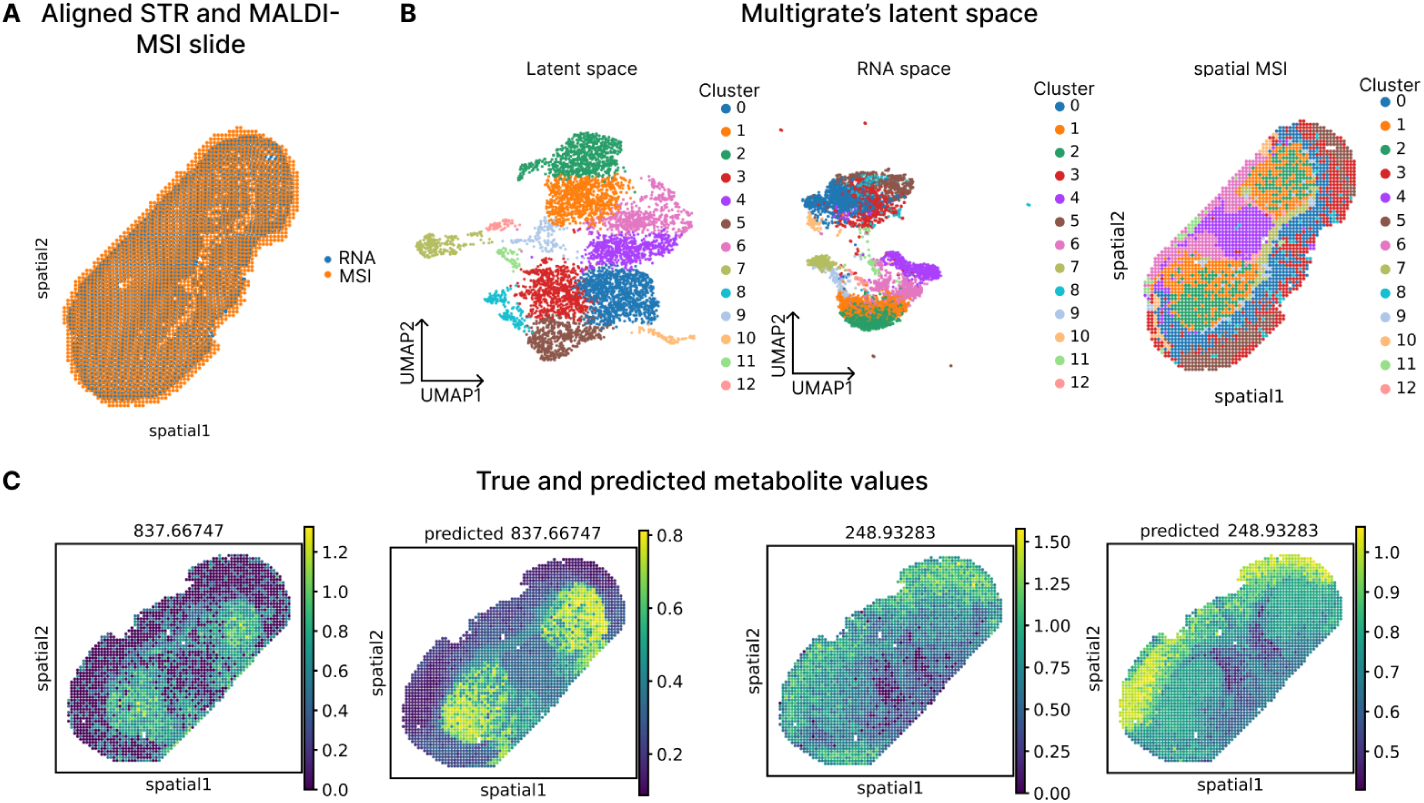
Integration and prediction on the MALDI-MSI data. **(A)** Spatial representation of the aligned spots from STR and MALDI-MSI. **(B)** UMAP of Multigrate’s integrated latent space colored by Leiden clusters; UMAP of the RNA measurements colored by Leiden clusters calculated on Multigrate’s latent space; spatial representation of MALDI-MSI spots colored by Leiden clusters calculated on Multigrate’s latent space. **(C)** True and predicted values for two metabolomic features exhibiting different spatial expression patterns.

## 4 Discussion

In this work, we introduced Multigrate, a scalable deep-learning model designed to integrate and query multimodal single-cell data. Our model utilized the product-of-experts approach and supports the integration of paired, partially overlapping, enabling query-to-reference mapping. One key advantage of Multigrate is its flexibility: new data modalities can be easily incorporated without the need for new model implementations, allowing for rapid exploratory analysis through a plug-and-play approach.

As a deep-learning method, Multigrate is subject to variability in downstream results due to the stochastic nature of the training process. Additionally, optimization of hyperparameters is necessary when new modalities or combinations of modalities are introduced.

We note that new metrics tailored specifically for multimodal integration are required to better assess the quality of the integrated latent space [30]. While some papers on multimodal integration use scIB metrics [8,36], others provide overviews of metrics explicitly introduced for the multimodal case [5]. Developing and standardizing such metrics will be crucial for future research.

Future work could explore using Jeffreys divergence, a symmetric version of KL divergence, to align distributions, as suggested in the MultiVI framework [4]. Additionally, as spatial metabolomics is still emerging, improvements in feature selection and data processing will enhance the prediction of metabolites, further expanding Multigrate’s capabilities for biological discovery. The implementation of the model is available at https://github.com/theislab/multigrate. The code necessary to reproduce the results is available at https://github.com/theislab/ multigrate_reproducibility.

## Acknowledgments

We thank Christopher Lance for the help with the cell type harmonization of the NeurIPS 2021 datasets, Michaela Müller for patiently answering all the pipeline infrastructure questions, Dr. Luke Zappia and Janneke Hulsen for figure feedback, and Dr. Malte Lücken for valuable feedback throughout the project. We thank the scverse community (especially the developers and the maintainers of `scanpy`, `muon` and `scvi-tools` packages) and all the Theislab for valuable discussions.

## Author contributions

A.L. and M.L. conceived the project with contributions from F.T. A.L. and M.L. designed the algorithm. A.L. implemented the algorithm with contributions from A.O. A.L. performed the paired integration, query-to-reference mapping and imputation experiments. A.L., M.S. and A.S. performed the mosaic integration and query-to-reference mapping experiments. F.C. curated the datasets and performed label harmonization for the first experiments in the project. M.S. implemented and performed the slide alignment for the spatial dataset. All authors contributed to the manuscript. M.L. and F.T. supervised the project.

## Supplementary Methods

### PoE-VAE model

We assume there are several single-cell multimodal datasets. Single-cell datasets are often confounded by the technical batch effect, but to simplify the notation, we will treat each dataset as one technical batch. In this section, we will refer to the experimental batches in an experiment or a dataset as “technical batches” or “batch covariates”. In contrast, the computational batches, i.e., mini-batches on which machine-learning models are trained, are referred to as “batches” or “training batches”.

We denote single-cell datasets as {*D*_1_*, …, D_k_*} with corresponding batch covariate labels {*c*_1_*, …, c_k_*} and assume that the datasets consist of patients {*p*_1_*, …, p_d_*} with corresponding disease labels {*l*_1_*, …, l_d_*}. We also assume that the datasets are multimodal and have *m* modalities in total.

We will now focus on a single mini-batch and describe one forward pass of the model. Each training batch consists of single-cell data {*X*_1_*, …, X_m_*}, the technical batch label {*c*}. The dataset information *c* is represented as a learnable embedding in a low-dimensional space. Hence, the batch input data matrices {*X*_1_*, …, X_m_*} correspond to multimodal data from one mini-batch from *m* modalities, where some matrices may be all zeros if measurements for the corresponding modality are missing. The number of rows in each matrix *X_i_* equals *n*, which is the number of cells in the mini-batch, and the number of columns equals the number of features in the original input data of modality *i*. Note that since the data is paired, the rows in different matrices within one batch always correspond to the same cells.

### Integration metrics

To assess the quality of the integration, we used several metrics from the scIB package [28]. Note that scIB metrics were designed for unimodal integration, and not all of them can be easily applied in the multimodal case; hence, we chose the metrics that only require the integrated embedding space as input (and not, e.g., the original unintegrated space). In the following, we briefly discuss two metrics for batch removal and four for biological variance conservation. As in scIB, the overall score was calculated as 0.4*batch correction + 0.6*biological conservation. For more details on the metrics and the implementation, see [28].

#### Batch correction

*Graph connectivity* measures how well cells from each cell type are connected in a k-nearest neighbor graph. If the connectivity is high, the batch effect was removed sufficiently. Average silhouette width (ASW) compares average distances within a cluster with distances to other clusters. The resulting score reflects how compact the clustering is. For *ASW batch*, we expect the batch clusters to be well-mixed together for a high batch correction score.

#### Biological variance conservation

*Adjusted Rand Index (ARI)* and *Normal-ized Mutual Information (NMI)* evaluate how well the clustering is aligned with the ground truth labels, i.e., cell type annotations. *ASW label* is a modification of the ASW batch, where we expect the cell type clusters to be compact and separate from other cell type clusters for a high biological conservation score. *Isolated label ASW* assesses how well rare cell types are distinguishable from the rest of the data.

### Benchmarks

#### Paired integration

To find the optimal hyperparameters for Multigrate, we performed a random grid search for the following parameters and values (with a maximum number of 100 iterations):

**Table 1.** Hyperparameter grid search for Multigrate’s paired integration.

| Hyperparameter | Description | Default | Range |
| --- | --- | --- | --- |
| Batch size | size of the training mini-batch | 256 | {128, 256, 512} |
| Learning rate | learning rate parameter | 1e-3 | {$1e-6$, $1e-5$, $1e-4$, $1e-3$} |
| KL coefficient | weight of KL loss in the overall loss | 1e-5 | {$1e-5$, $1e-4$, $1e-3$, $1e-2$, $1e-1$} |
| Latent dimension | dimensionality of the latent space | 16 | {8, 16, 32} |
| Conditional dimension | dimensionality of the covariate embedding space | 16 | {8, 16, 32} |
| Number of layers | number of hidden layers in encoders and decoders | 1 | {0, 1, 2} |
| Activation function | non-linearity in the network | LeakyReLU | {LeakyReLU, Tanh} |

#### Integration for query-to-reference mapping and prediction of missing modalities

We performed a hyperparameter search for PoE and MoE models for the following parameters and values:

**Table 2.** Hyperparameter search for Multigrate’s PoE vs MoE.

| Hyperparameter | Description | Range |
| --- | --- | --- |
| Learning rate | learning rate parameter | {$1e-4$, $1e-3$} |
| KL coefficient | weight of KL loss in the overall loss | {$1e-5$, $1e-4$} |
| Integration coefficient | weight of integration MMD loss in the overall loss | {1, 10, 100, $1e3$, $1e4$, $1e5$} |
| Modality 1 loss | weight for the modality 1 loss | {0.1, 1, 10} |
| Modality 2 loss | weight for the modality 2 loss | {0.1, 1, 10} |
| MMD loss | type of the MMD loss | {‘latent’, ‘marginal’} |

The rest of the parameters were set to the default values from Table 1.

#### Mosaic query-to-reference mapping

Seurat v5 and Multigrate allow query-to-reference mapping onto the atlases. For Seurat’s bridge integration, we first build an RNA-seq-only reference atlas from scRNA-seq measurements from the CITE-seq dataset and snRNA-seq measurements from the multiome dataset using data from Site 1 and Site 2. Then we used one donor (donor 7) from Site 3 as a CITE-seq bridge to map protein data from Site 4 (donor 9) on top of the RNA-seq reference and the same donor from Site 3 as a multiome bridge to map scATAC-seq data from Site 4 (donor 9) onto the same reference.

For Seurat’s Bridge, the reference is a scRNA-seq-only reference (i.e., not multimodal), so we could not directly compare the reference building with the other methods for the mosaic reference building. Additionally, Bridge allows visualization of the reference and query on a joint UMAP and label transfer but does not explicitly provide low-dimensional embeddings in the joint referencequery space. Hence, we did not calculate scIB metrics for Seurat Bridge, but we included UMAPs of the reference and the mapped scATAC-seq query in the supplementary figures for visual inspection (Suppl. Fig. 5A,B)

We performed a hyperparameter search for Multigrate for the following parameters and values:

**Table 3.**
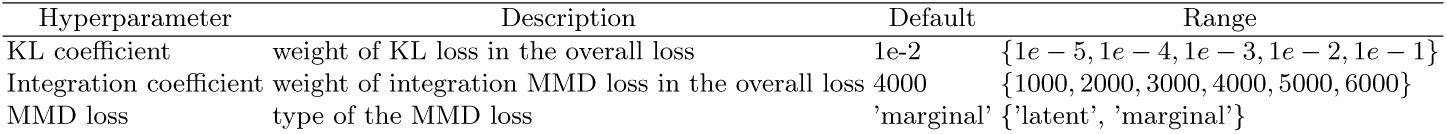
Hyperparameter search for Multigrate’s mosaic integration and query-to-reference mapping.

MMD loss type refers to how we calculate the MMD loss: ‘latent’ means that 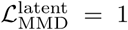 and 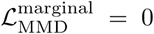; ‘marginal’ means that 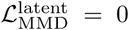 and 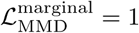.

Other hyperparameters were set to their defaults from Table 1. To choose the default parameters, we calculated the scIB metrics on the reference and the mapped queries (with the batch covariate indicating whether the cell came from the reference or the query) to assess the mapping quality.

To assess the accuracy of cell-type transfer, we trained random forest classifiers for each of the query types with `sklearn.ensemble.RandomForestClassifier()` specifying the parameter `class_weight` as “balanced_subsample”.

#### Robustness of the integration module

To assess the robustness of the integration, we performed several experiments on the mosaic dataset. We tested several parameters: integration coefficient (i.e., MMD coefficient *λ*_MMD_), number of shared features between datasets from different technologies, selection of integration covariates, reference/query ratio and different ways of calculating the MMD loss. Unless the parameter was tested in the experiment, the default parameters used throughout this benchmark were taken from Table 1, and the rest is shown in Table 4.

### Default architecture

The integration module consists of encoder-decoder pairs, and below we provide the specifications of each pair. Mu and Sigma modules output the *µ* and *σ* parameters of the unimodal distributions. Unless specified, the parameters have their default values from PyTorch.

**Table 4.** Parameters tested in the robustness benchmark.

| Hyperparameter | Description | Default | Range |
| --- | --- | --- | --- |
| Integration coefficient | weight of the MMD loss in the overall loss | 1e4 | {1e-3, 1e-2, 1e-1, 1, 10, 1e2, 1e3, 1e4, 1e5, 1e6, 1e7} |
| Number of shared features | number of shared features between scRNA and snRNA | 4000 | {100, 500, 1000, 2000, 3000, 4000} |
| Integration covariate | covariate used for the calculation of MMD | modality | {none, modality, donor} |
| Batch covariate | covariate(s) used as batch covariate(s) | modality & donor | {modality, donor, modality & donor} |
| Reference/query split | which sites were used as reference and which as query | Sites 1-3/Site 4 | {Sites 1-3/Site 4, Sites 1-2/Sites 3-4, Site 1/Sites 2-4} |
| MMD type | how MMD loss was calculated | marginal | {marginal, latent} |

**Table 5.**
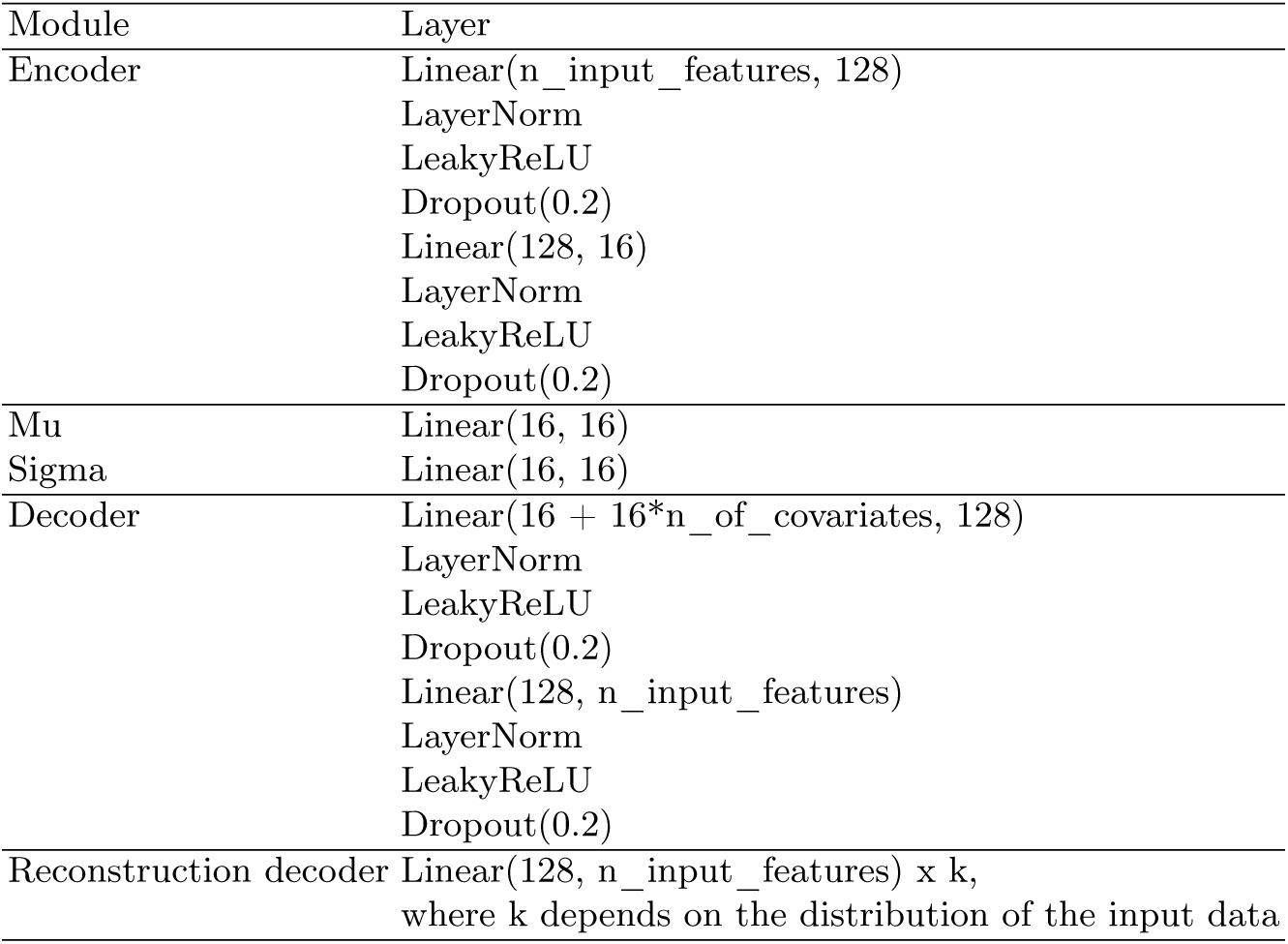

| Module | Layer |
| --- | --- |
| Encoder | Linear(n_input_features, 128) |
|  | LayerNorm |
|  | LeakyReLU |
|  | Dropout(0.2) |
|  | Linear(128, 16) |
|  | LayerNorm |
|  | LeakyReLU |
|  | Dropout(0.2) |
| Mu | Linear(16, 16) |
| Sigma | Linear(16, 16) |
| Decoder | Linear(16 + 16*n_of_covariates, 128) |
|  | LayerNorm |
|  | LeakyReLU |
|  | Dropout(0.2) |
|  | Linear(128, n_input_features) |
|  | LayerNorm |
|  | LeakyReLU |
|  | Dropout(0.2) |
| Reconstruction decoder | Linear(128, n_input_features) x k,<br>where k depends on the distribution of the input data |

**Supplementary Figure 1.**
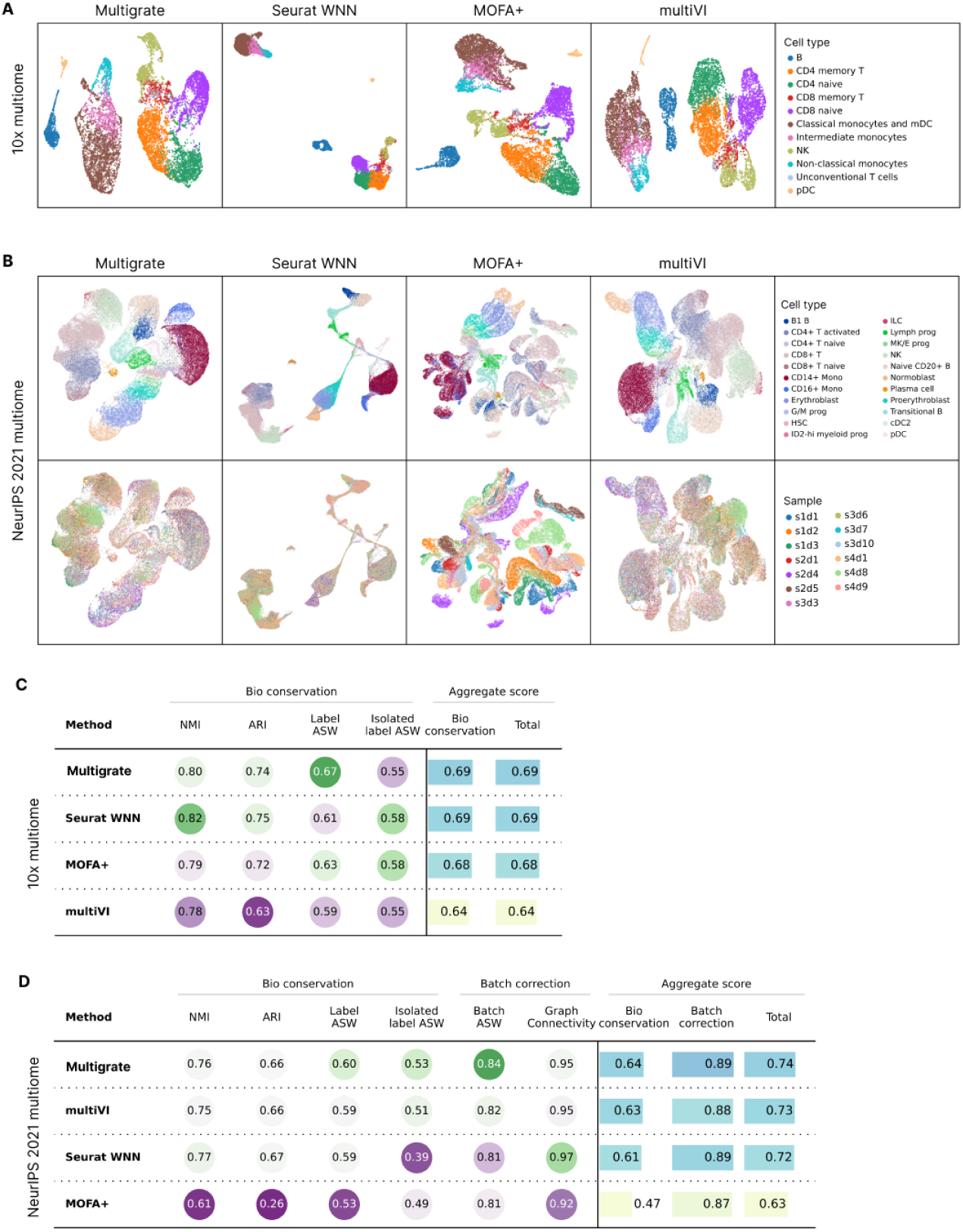
Paired integration of multiome datasets. **(A)** UMAPs of the latent spaces of the 10x multiome dataset, integrated with Multigrate, Seurat WNN, MOFA+ and multiVI, colored by cell type. **(B)** UMAPs of the latent spaces of the NeurIPS 2021 multiome dataset, integrated with Multigrate, Seurat WNN, MOFA+ and multiVI, colored by cell type and sample. **(C)** A table showing scIB metric scores for 10x multiome dataset. **(D)** A table showing scIB metric scores for NeurIPS 2021 multiome dataset.

**Supplementary Figure 2.**
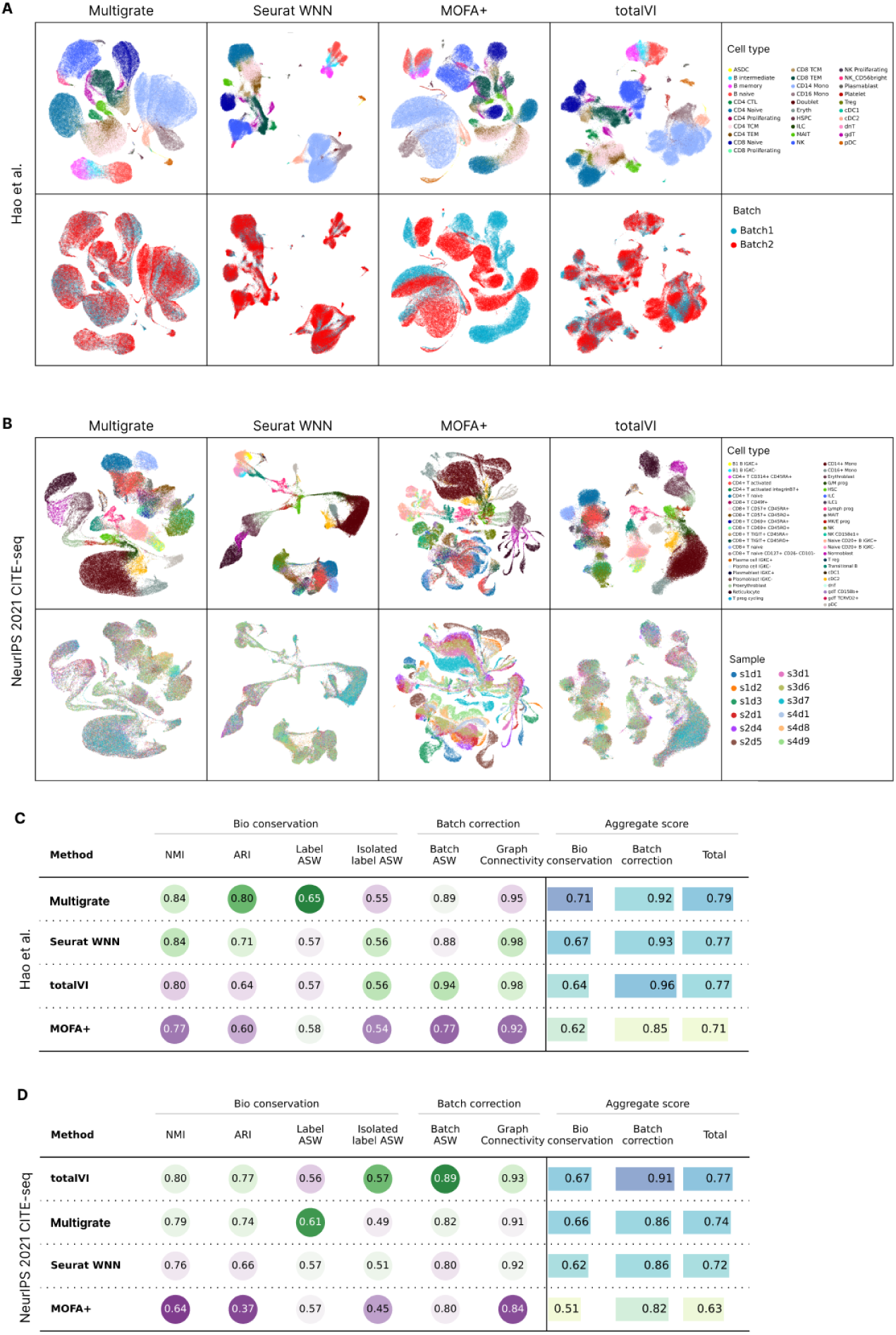
Paired integration of CITE-seq datasets. **(A)** UMAPs of the latent spaces of the Hao *el at.* dataset, integrated with Multigrate, Seurat WNN, MOFA+ and totalVI, colored by cell type and batch. **(B)** UMAPs of the latent spaces of the NeurIPS 2021 CITE-seq dataset, integrated with Multigrate, Seurat WNN, MOFA+ and totalVI, colored by cell type and sample. **(C)** A table showing scIB metric scores for Hao *et al.* CITE-seq dataset. **(D)** A table showing scIB metric scores for NeurIPS 2021 CITE-seq dataset.

**Supplementary Figure 3.**
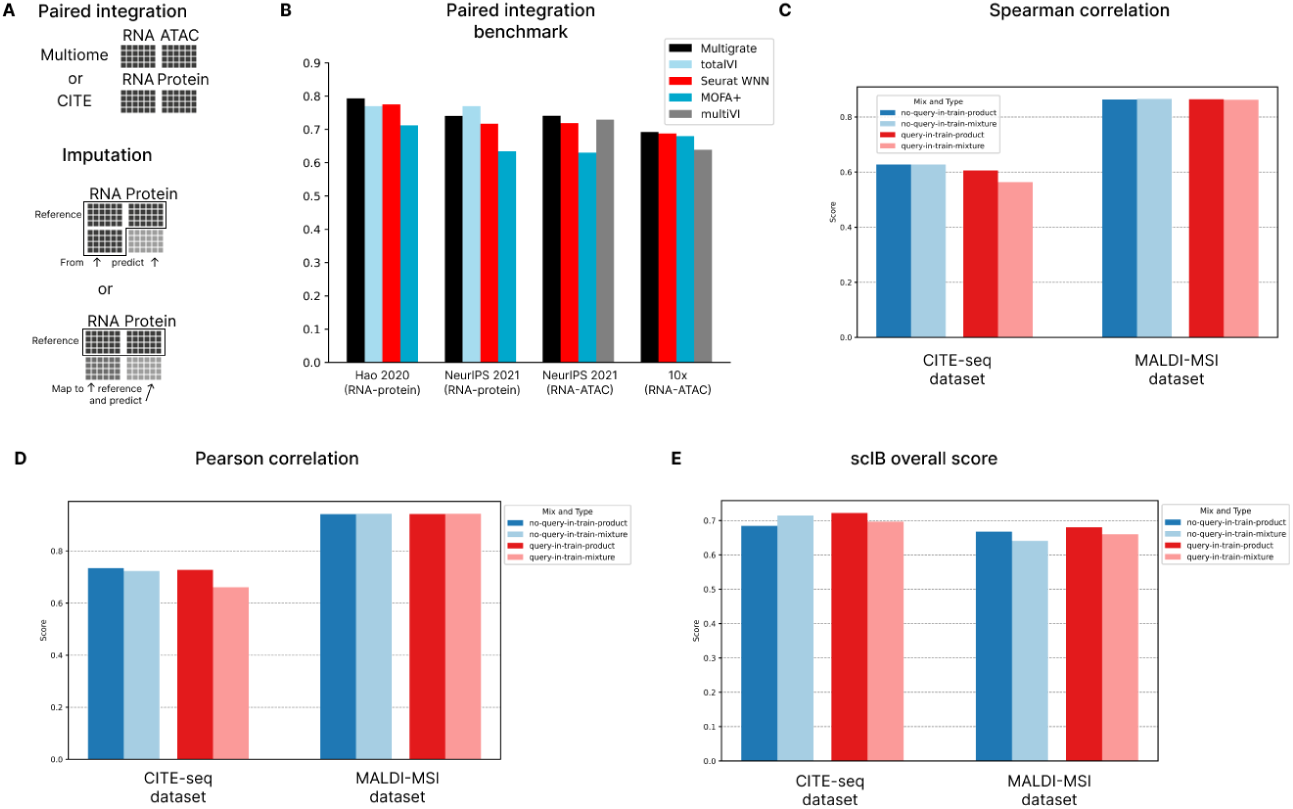
Overall scores for paired integration and query-to-reference mapping. **(A)** Paired integration was performed for CITE-seq (RNA-protein) and multiome (RNA-ATAC) datasets, where observations, i.e., cells, are matched. For the imputation experiments, the reference was either a paired dataset with another RNA-only batch or only the paired dataset. In the former case, the missing modality was predicted directly after training for the RNA-only batch. In the latter case, a new RNA-only batch was mapped onto the reference with scArches and then the missing modality was predicted. **(B)** Overall scIB scores for 2 CITE-seq datasets and 2 multiome datasets reported for Multigrate, Seurat WNN, MOFA+, totalVI for CITE-seq data and multiVI for multiome data. **(C)** Spearman correlation between true and predicted values for the missing modality imputation experiments. Values in **(C-E)** are reported for NeurIPS 2021 CITE-seq and Vicari et al. datasets for PoE and MoE models in both query-mapping settings. **(D)** Pearson correlation between true and predicted values for the missing modality imputation experiments. **(D)** Overall scIB score the missing modality imputation experiments. The metrics assess the quality of the query mapping.

**Supplementary Figure 4.**
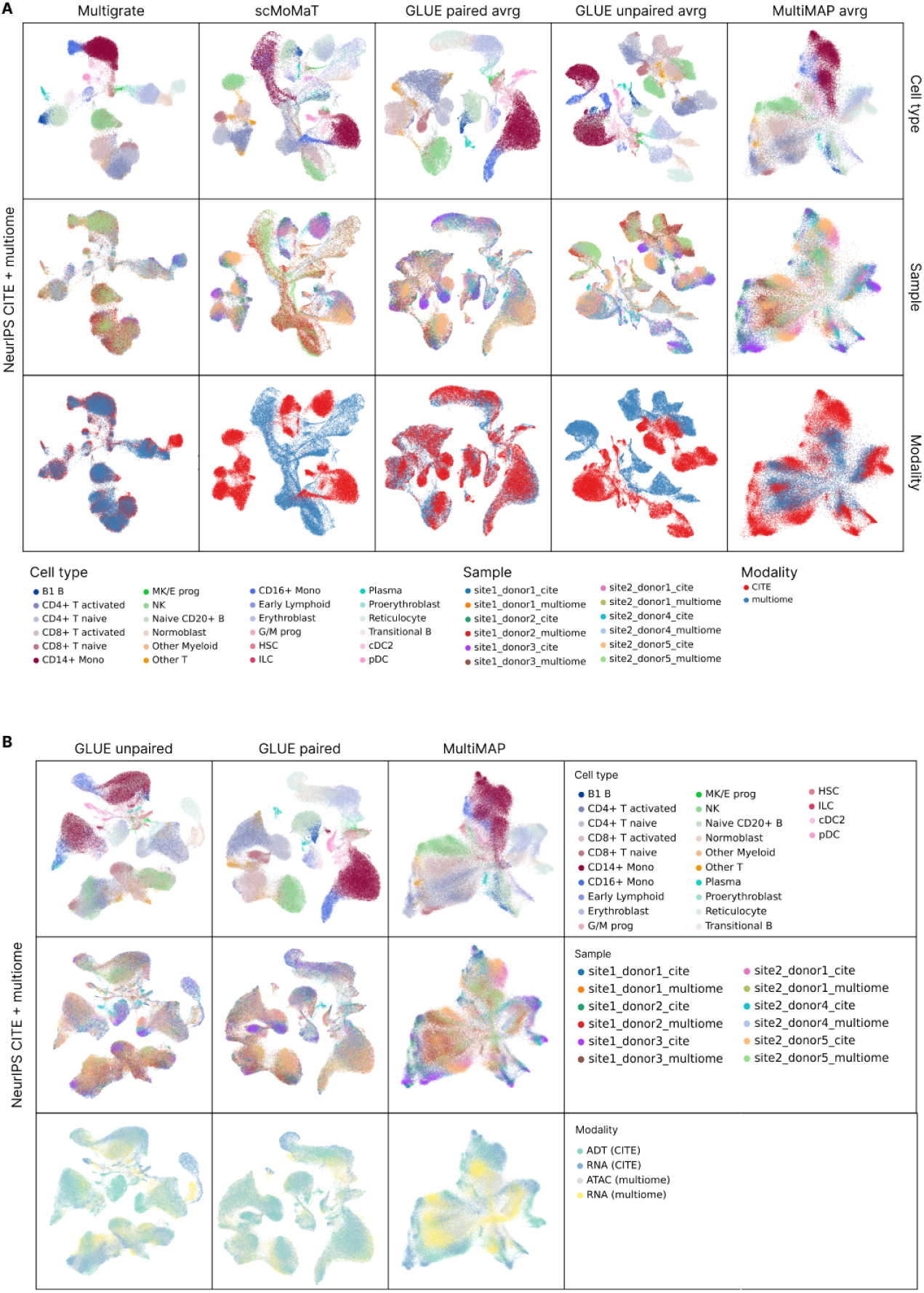
Trimodal reference building. **(a)** UMAPs of the latent spaces of NeurIPS 2021 multiome and NeurIPS 2021 CITE-seq datasets, integrated with methods that output a representation per cell, i.e., Multigrate, scMoMaT, GLUE paired (averaged representation), GLUE unpaired (averaged representation) and MultiMAP (averaged representation), colored by cell type, sample and modality. **(b)** UMAPs of the latent spaces of NeurIPS 2021 multiome and NeurIPS 2021 CITE-seq datasets, integrated with methods that output a representation per cell per modality, i.e., GLUE unpaired, GLUE paired and MultiMAP, colored by cell type, sample and modality.

**Supplementary Figure 5.**
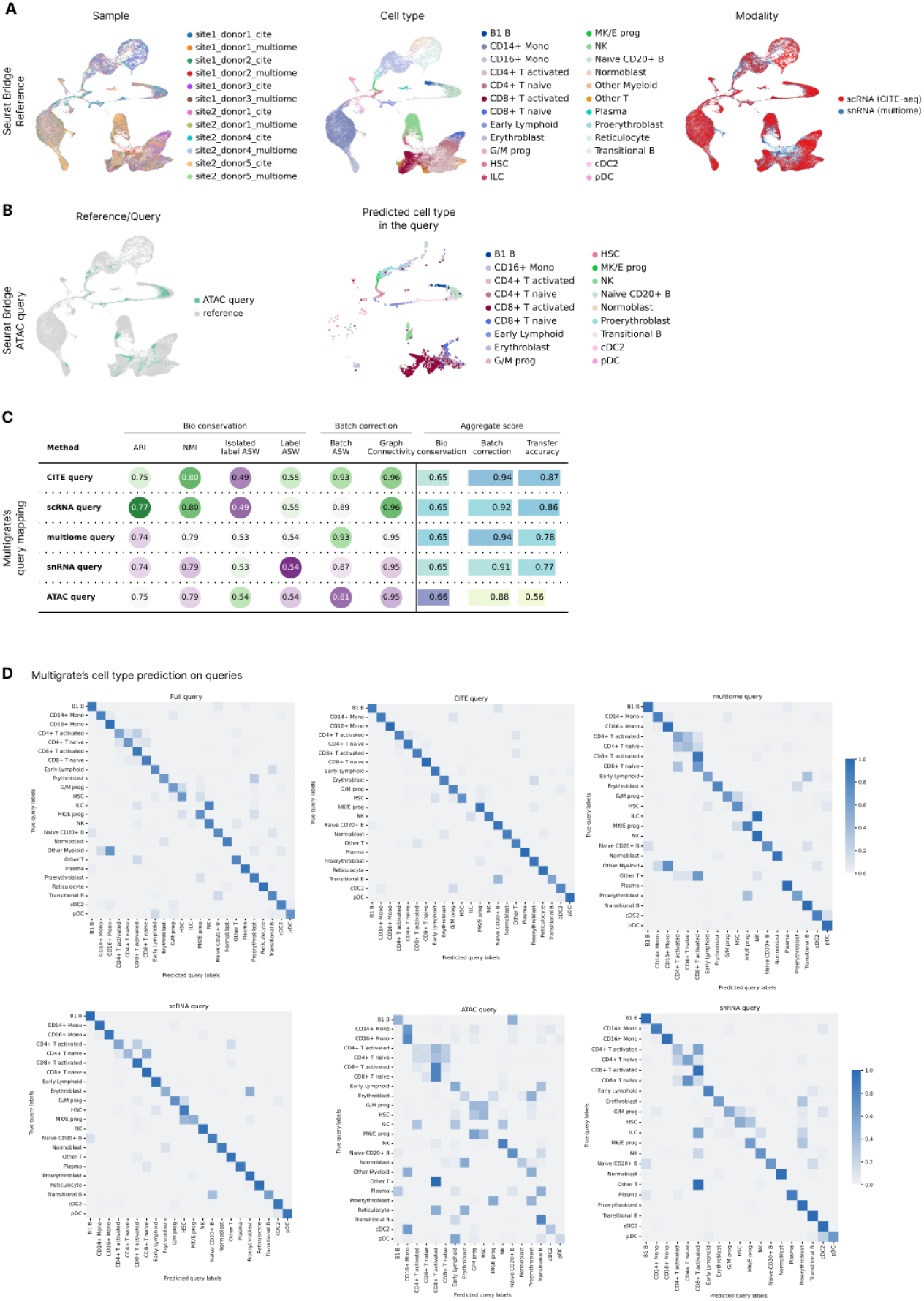
Trimodal query mapping. **(a)** UMAPs of the integrated scRNA-seq and snRNA-seq from NeurIPS 2021 CITE-seq and NeurIPS 2021 multiome, respectively, with Seurat, colored by sample, cell type and modality/dataset. **(b)** UMAPs of the mapped ATAC query onto the RNA-seq reference with Bridge colored by reference/query and ATAC query only colored by cell type. **(c)** A table with scIB scores calculated for different queries mapped with Multigrate. **(d)** Confusion matrices between true and predicted (with a random forest model) cell types for the full query and individual queries mapped with Multigrate.

**Supplementary Figure 6.**
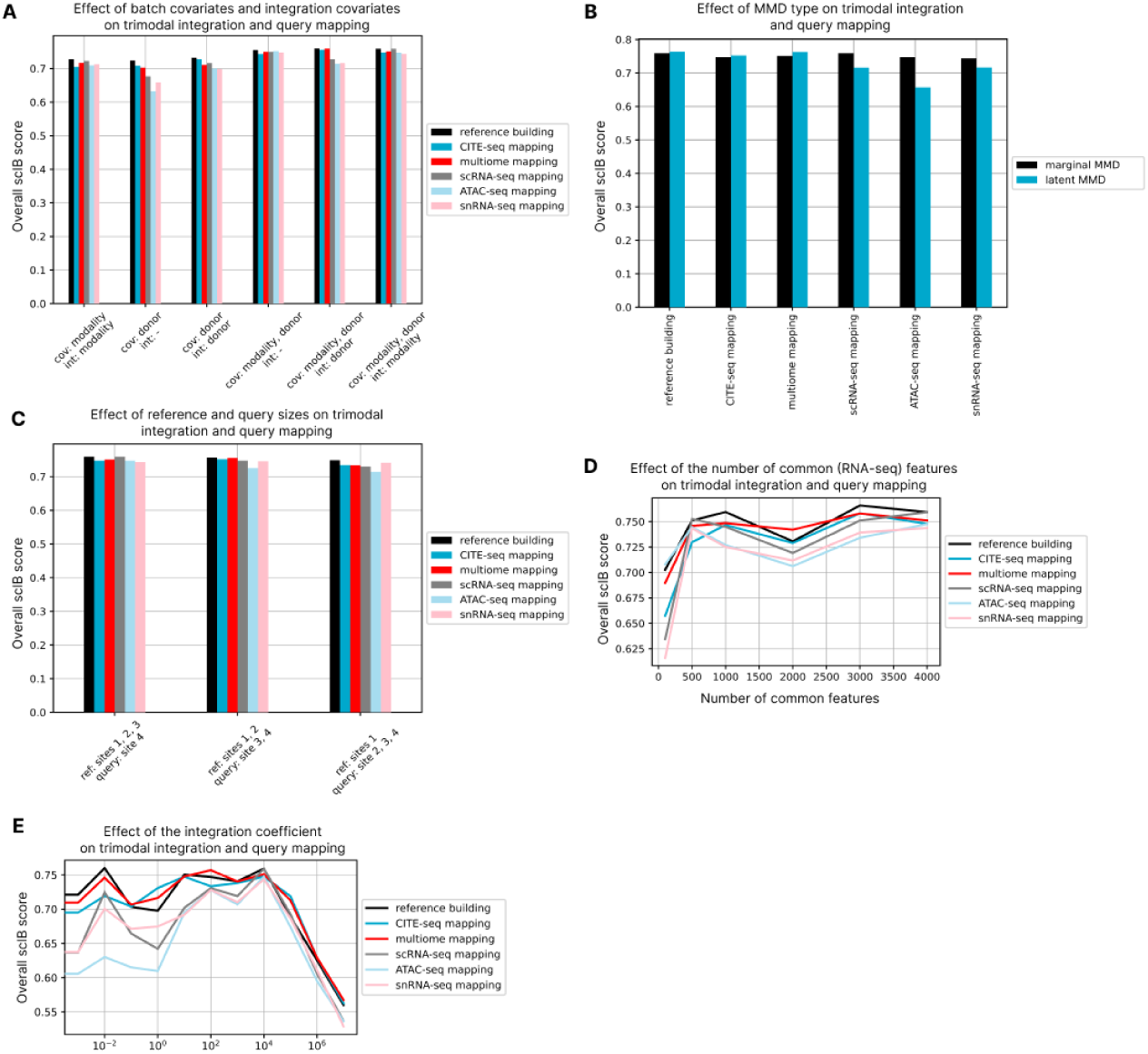
Robustness of trimodal integration with Multigrate. **(a)** A bar plot showing the effect of batch covariates and integration covariates selection on the scIB overall integration score and query mapping scores. **(b)** A bar plot showing the effect of MMD loss type on the scIB overall integration score and query mapping scores. **(c)** A bar plot showing the effect of the reference and query sizes on the scIB overall integration score and query mapping scores. **(d)** A line plot showing the effect of the number of the common features in the scRNA/snRNA modality on the scIB overall integration score and query mapping scores. **(e)** A line plot showing the effect of the integration coefficient (i.e., the weight of the MMD loss) on the scIB overall integration score and query mapping scores.

## References

1. Datasets - single cell multiome atac + gene exp. - official 10x genomics support. https://support.10xgenomics.com/single-cell-multiome-atac-gex/datasets/2.0.0/pbmc_granulocyte_sorted_10k

2. Argelaguet, R., Arnol, D., Bredikhin, D., Deloro, Y., Velten, B., Marioni, J.C., Stegle, O.: Mofa+: a statistical framework for comprehensive integration of multi-modal single-cell data. Genome Biology 21(1), 111 (May 2020). 10.1186/s13059-020-02015-1, 10.1186/s13059-020-02015-1

3. Argelaguet, R., Cuomo, A.S.E., Stegle, O., Marioni, J.C.: Computational principles and challenges in single-cell data integration. Nat. Biotechnol. 39(10), 1202–1215 (Oct 2021)

4. Ashuach, T., Gabitto, M.I., Koodli, R.V., Saldi, G.A., Jordan, M.I., Yosef, N.: MultiVI: deep generative model for the integration of multimodal data. Nat. Methods 20(8), 1222–1231 (Aug 2023)

5. Athaya, T., Ripan, R.C., Li, X., Hu, H.: Multimodal deep learning approaches for single-cell multi-omics data integration. Brief. Bioinform. (Aug 2023)

6. Barshan, E., Ghodsi, A., Azimifar, Z., Zolghadri Jahromi, M.: Supervised principal component analysis: Visualization, classification and regression on subspaces and submanifolds. Pattern Recognition 44(7), 1357–1371 (2011), https://www. sciencedirect.com/science/article/pii/S0031320310005819

7. Baysoy, A., Bai, Z., Satija, R., Fan, R.: The technological landscape and applications of single-cell multi-omics. Nat. Rev. Mol. Cell Biol. pp. 1–19 (Jun 2023)

8. Brombacher, E., Hackenberg, M., Kreutz, C., Binder, H., Treppner, M.: The performance of deep generative models for learning joint embeddings of single-cell multi-omics data. Front Mol Biosci 9, 962644 (Oct 2022)

9. Bunne, C., Roohani, Y., Rosen, Y., Gupta, A., Zhang, X., Roed, M., Alexandrov, T., AlQuraishi, M., Brennan, P., Burkhardt, D.B., Califano, A., Cool, J., Dernburg, A.F., Ewing, K., Fox, E.B., Haury, M., Herr, A.E., Horvitz, E., Hsu, P.D., Jain, V., Johnson, G.R., Kalil, T., Kelley, D.R., Kelley, S.O., Kreshuk, A., Mitchison, T., Otte, S., Shendure, J., Sofroniew, N.J., Theis, F., Theodoris, C.V., Upadhyayula, S., Valer, M., Wang, B., Xing, E., Yeung-Levy, S., Zitnik, M., Karaletsos, T., Regev, A., Lundberg, E., Leskovec, J., Quake, S.R.: How to build the virtual cell with artificial intelligence: Priorities and opportunities. arXiv [q-bio.QM] (Sep 2024)

10. Cao, Z.J., Gao, G.: Multi-omics single-cell data integration and regulatory inference with graph-linked embedding. Nat. Biotechnol. 40(10), 1458–1466 (Oct 2022)

11. Danese, A., Richter, M.L., Chaichoompu, K., Fischer, D.S., Theis, F.J., Colomé-Tatché, M.: EpiScanpy: integrated single-cell epigenomic analysis. Nat. Commun. 12(1), 5228 (Sep 2021)

12. Drost, F., An, Y., Bonafonte-Pardàs, I., Dratva, L.M., Lindeboom, R.G.H., Haniffa, M., Teichmann, S.A., Theis, F., Lotfollahi, M., Schubert, B.: Multi-modal generative modeling for joint analysis of single-cell T cell receptor and gene expression data. Nat. Commun. 15(1), 5577 (Jul 2024)

13. Gayoso, A., Steier, Z., Lopez, R., Regier, J., Nazor, K.L., Streets, A., Yosef, N.: Joint probabilistic modeling of single-cell multi-omic data with totalvi. Nature Methods 18(3), 272–282 (Mar 2021). 10.1038/s41592-020-01050-x, 10.1038/s41592-020-01050-x

14. Gretton, A., Borgwardt, K., Rasch, M., Schölkopf, B., Smola, A.: A kernel Two-Sample test. J. Mach. Learn. Res. 13, 723–773 (Mar 2012)

15. Hao, Y., Hao, S., Andersen-Nissen, E., Mauck, W.M., Zheng, S., Butler, A., Lee, M.J., Wilk, A.J., Darby, C., Zager, M., Hoffman, P., Stoeckius, M., Papalexi, E., Mimitou, E.P., Jain, J., Srivastava, A., Stuart, T., Fleming, L.M., Yeung, B., Rogers, A.J., McElrath, J.M., Blish, C.A., Gottardo, R., Smibert, P., Satija, R.: Integrated analysis of multimodal single-cell data. Cell 184(13), 3573–3587.e29 (2021). 10.1016/j.cell.2021.04.048, https://www.sciencedirect.com/science/article/pii/S0092867421005833

16. Hinton, G.E.: Training products of experts by minimizing contrastive divergence. Neural Comput. 14(8), 1771–1800 (Aug 2002)

17. Jain, M.S., Polanski, K., Conde, C.D., Chen, X., Park, J., Mamanova, L., Knights, A., Botting, R.A., Stephenson, E., Haniffa, M., Lamacraft, A., Efremova, M., Teichmann, S.A.: MultiMAP: dimensionality reduction and integration of multimodal data. Genome Biol. 22(1), 346 (Dec 2021)

18. Kaya-Okur, H.S., Wu, S.J., Codomo, C.A., Pledger, E.S., Bryson, T.D., Henikoff, J.G., Ahmad, K., Henikoff, S.: CUT&Tag for efficient epigenomic profiling of small samples and single cells. Nat. Commun. 10(1), 1930 (Apr 2019)

19. Kingma, D.P., Salimans, T., Welling, M.: Variational dropout and the local reparameterization trick. In: Cortes, C., Lawrence, N., Lee, D., Sugiyama, M., Garnett, R. (eds.) Advances in Neural Information Processing Systems. vol. 28. Curran Associates, Inc. (2015), https://proceedings.neurips.cc/paper_files/paper/2015/file/bc7316929fe1545bf0b98d114ee3ecb8-Paper.pdf

20. Klein, D., Palla, G., Lange, M., Klein, M., Piran, Z., Gander, M., Meng-Papaxanthos, L., Sterr, M., Bastidas-Ponce, A., Tarquis-Medina, M., Lickert, H., Bakhti, M., Nitzan, M., Cuturi, M., Theis, F.J.: Mapping cells through time and space with moscot. Bioinformatics (biorxiv;2023.05.11.540374v2) (May 2023)

21. Lance, C., Luecken, M.D., Burkhardt, D.B., Cannoodt, R., Rautenstrauch, P., Laddach, A., Ubingazhibov, A., Cao, Z.J., Deng, K., Khan, S., Liu, Q., Russkikh, N., Ryazantsev, G., Ohler, U., data integration competition participants, N..M., Pisco, A.O., Bloom, J., Krishnaswamy, S., Theis, F.J.: Multimodal single cell data integration challenge: Results and lessons learned. In: Kiela, D., Ciccone, M., Caputo, B. (eds.) Proceedings of the NeurIPS 2021 Competitions and Demonstrations Track. Proceedings of Machine Learning Research, vol. 176, pp. 162–176. PMLR (06–14 Dec 2022), https://proceedings.mlr.press/v176/lance22a.html

22. Lee, C., van der Schaar, M.: A variational information bottleneck approach to multi-omics data integration. In: Proceedings of The 24th International Conference on Artificial Intelligence and Statistics. Proceedings of Machine Learning Research, vol. 130, pp. 1513–1521 (13–15 Apr 2021), http://proceedings.mlr.press/v130/lee21a.html

23. Li, J., Wang, J., Ibarra, I.L., Cheng, X., Luecken, M.D., Lu, J., Monavarfeshani, A., Yan, W., Zheng, Y., Zuo, Z., Colborn, S.L.Z., Cortez, B.S., Owen, L.A., Tran, N.M., Shekhar, K., Sanes, J.R., Timothy Stout, J., Chen, S., Li, Y., DeAngelis, M.M., Theis, F.J., Chen, R.: Integrated multi-omics single cell atlas of the human retina. bioRxiv p. 2023.11.07.566105 (Nov 2023)

24. Lotfollahi, M., Litinetskaya, A., Theis, F.J.: Multigrate: single-cell multi-omic data integration. bioRxiv p. 2022.03.16.484643 (Mar 2022)

25. Lotfollahi, M., Naghipourfar, M., Luecken, M.D., Khajavi, M., Büttner, M., Wagenstetter, M., Avsec, Ž., Gayoso, A., Yosef, N., Interlandi, M., Rybakov, S., Misharin, A.V., Theis, F.J.: Mapping single-cell data to reference atlases by transfer learning. Nat. Biotechnol. 40(1), 121–130 (Jan 2022)

26. Lotfollahi, M., Naghipourfar, M., Theis, F.J., Wolf, F.A.: Conditional out-of-distribution generation for unpaired data using transfer VAE. Bioinformatics 36(Suppl_2), i610–i617 (Dec 2020)

27. Luecken, M., Burkhardt, D., Cannoodt, R., Lance, C., Agrawal, A., Aliee, H., Chen, A., Deconinck, L., Detweiler, A., Granados, A., Huynh, S., Isacco, L., Kim, Y., Klein, D., De Kumar, B., Kuppasani, S., Lickert, H., McGeever, A., Melgarejo, J., Mekonen, H., Morri, M., Müller, M., Neff, N., Paul, S., Rieck, B., Schneider, K., Steelman, S., Sterr, M., Treacy, D., Tong, A., Villani, A.C., Wang, G., Yan, J., Zhang, C., Pisco, A., Krishnaswamy, S., Theis, F., Bloom, J.M.: A sandbox for prediction and integration of dna, rna, and proteins in single cells. In: Vanschoren, J., Yeung, S. (eds.) Proceedings of the Neural Information Processing Systems Track on Datasets and Benchmarks. vol. 1. Curran (2021), https://datasets-benchmarks-proceedings.neurips.cc/paper_files/paper/2021/file/158f3069a435b314a80bdcb024f8e422-Paper-round2.pdf

28. Luecken, M.D., Büttner, M., Chaichoompu, K., Danese, A., Interlandi, M., Mueller, M.F., Strobl, D.C., Zappia, L., Dugas, M., Colomé-Tatché, M., Theis, F.J.: Benchmarking atlas-level data integration in single-cell genomics. Nat. Methods 19(1), 41–50 (Jan 2022)

29. Minoura, K., Abe, K., Nam, H., Nishikawa, H., Shimamura, T.: A mixture-of-experts deep generative model for integrated analysis of single-cell multiomics data. Cell Rep. Methods 1(5), 100071 (Sep 2021)

30. Rautenstrauch, P., Vlot, A.H.C., Saran, S., Ohler, U.: Intricacies of single-cell multi-omics data integration. Trends Genet. 38(2), 128–139 (Feb 2022)

31. Sikkema, L., Ramírez-Suástegui, C., Strobl, D.C., Gillett, T.E., Zappia, L., Madissoon, E., Markov, N.S., Zaragosi, L.E., Ji, Y., Ansari, M., Arguel, M.J., Apperloo, L., Banchero, M., Bécavin, C., Berg, M., Chichelnitskiy, E., Chung, M.I., Collin, A., Gay, A.C.A., Gote-Schniering, J., Hooshiar Kashani, B., Inecik, K., Jain, M., Kapellos, T.S., Kole, T.M., Leroy, S., Mayr, C.H., Oliver, A.J., von Papen, M., Peter, L., Taylor, C.J., Walzthoeni, T., Xu, C., Bui, L.T., De Donno, C., Dony, L., Faiz, A., Guo, M., Gutierrez, A.J., Heumos, L., Huang, N., Ibarra, I.L., Jackson, N.D., Kadur Lakshminarasimha Murthy, P., Lotfollahi, M., Tabib, T., Talavera-López, C., Travaglini, K.J., Wilbrey-Clark, A., Worlock, K.B., Yoshida, M., Lung Biological Network Consortium, van den Berge, M., Bossé, Y., Desai, T.J., Eickelberg, O., Kaminski, N., Krasnow, M.A., Lafyatis, R., Nikolic, M.Z., Powell, J.E., Rajagopal, J., Rojas, M., Rozenblatt-Rosen, O., Seibold, M.A., Sheppard, D., Shepherd, D.P., Sin, D.D., Timens, W., Tsankov, A.M., Whitsett, J., Xu, Y., Banovich, N.E., Barbry, P., Duong, T.E., Falk, C.S., Meyer, K.B., Kropski, J.A., Pe’er, D., Schiller, H.B., Tata, P.R., Schultze, J.L., Teichmann, S.A., Misharin, A.V., Nawijn, M.C., Luecken, M.D., Theis, F.J.: An integrated cell atlas of the lung in health and disease. Nat. Med. 29(6), 1563–1577 (Jun 2023)

32. Sohn, K., Lee, H., Yan, X.: Learning structured output representation using deep conditional generative models. In: Cortes, C., Lawrence, N., Lee, D., Sugiyama, M., Garnett, R. (eds.) Advances in Neural Information Processing Systems. vol. 28. Curran Associates, Inc. (2015), https://proceedings.neurips.cc/paper_files/paper/2015/file/8d55a249e6baa5c06772297520da2051-Paper.pdf

33. Stoeckius, M., Hafemeister, C., Stephenson, W., Houck-Loomis, B., Chattopadhyay, P.K., Swerdlow, H., Satija, R., Smibert, P.: Simultaneous epitope and transcriptome measurement in single cells. Nature Methods 14(9), 865–868 (Sep 2017), 10.1038/nmeth.4380

34. Traag, V.A., Waltman, L., van Eck, N.J.: From louvain to leiden: guaranteeing well-connected communities. Sci. Rep. 9(1), 5233 (Mar 2019)

35. Vicari, M., Mirzazadeh, R., Nilsson, A., Shariatgorji, R., Bjärterot, P., Larsson, L., Lee, H., Nilsson, M., Foyer, J., Ekvall, M., Czarnewski, P., Zhang, X., Svenningsson, P., Käll, L., Andrén, P.E., Lundeberg, J.: Spatial multimodal analysis of transcriptomes and metabolomes in tissues. Nat. Biotechnol. 42(7), 1046–1050 (Jul 2024)

36. Xiao, C., Chen, Y., Meng, Q., Wei, L., Zhang, X.: Benchmarking multi-omics integration algorithms across single-cell RNA and ATAC data. Briefings in Bioinformatics 25(2), bbae095 (03 2024). 10.1093/bib/bbae095, 10.1093/bib/bbae095

37. Yao, Z., Liu, H., Xie, F., Fischer, S., Adkins, R.S., Aldridge, A.I., Ament, S.A., Bartlett, A., Behrens, M.M., Van den Berge, K., Bertagnolli, D., de Bézieux, H.R., Biancalani, T., Booeshaghi, A.S., Bravo, H.C., Casper, T., Colantuoni, C., Crabtree, J., Creasy, H., Crichton, K., Crow, M., Dee, N., Dougherty, E.L., Doyle, W.I., Dudoit, S., Fang, R., Felix, V., Fong, O., Giglio, M., Goldy, J., Hawrylycz, M., Herb, B.R., Hertzano, R., Hou, X., Hu, Q., Kancherla, J., Kroll, M., Lathia, K., Li, Y.E., Lucero, J.D., Luo, C., Mahurkar, A., McMillen, D., Nadaf, N.M., Nery, J.R., Nguyen, T.N., Niu, S.Y., Ntranos, V., Orvis, J., Osteen, J.K., Pham, T., Pinto-Duarte, A., Poirion, O., Preissl, S., Purdom, E., Rimorin, C., Risso, D., Rivkin, A.C., Smith, K., Street, K., Sulc, J., Svensson, V., Tieu, M., Torkelson, A., Tung, H., Vaishnav, E.D., Vanderburg, C.R., van Velthoven, C., Wang, X., White, O.R., Huang, Z.J., Kharchenko, P.V., Pachter, L., Ngai, J., Regev, A., Tasic, B., Welch, J.D., Gillis, J., Macosko, E.Z., Ren, B., Ecker, J.R., Zeng, H., Mukamel, E.A.: A transcriptomic and epigenomic cell atlas of the mouse primary motor cortex. Nature 598(7879), 103–110 (Oct 2021)

38. Yao, Z., van Velthoven, C.T.J., Kunst, M., Zhang, M., McMillen, D., Lee, C., Jung, W., Goldy, J., Abdelhak, A., Aitken, M., Baker, K., Baker, P., Barkan, E., Bertagnolli, D., Bhandiwad, A., Bielstein, C., Bishwakarma, P., Campos, J., Carey, D., Casper, T., Chakka, A.B., Chakrabarty, R., Chavan, S., Chen, M., Clark, M., Close, J., Crichton, K., Daniel, S., DiValentin, P., Dolbeare, T., Ellingwood, L., Fiabane, E., Fliss, T., Gee, J., Gerstenberger, J., Glandon, A., Gloe, J., Gould, J., Gray, J., Guilford, N., Guzman, J., Hirschstein, D., Ho, W., Hooper, M., Huang, M., Hupp, M., Jin, K., Kroll, M., Lathia, K., Leon, A., Li, S., Long, B., Madigan, Z., Malloy, J., Malone, J., Maltzer, Z., Martin, N., McCue, R., McGinty, R., Mei, N., Melchor, J., Meyerdierks, E., Mollenkopf, T., Moonsman, S., Nguyen, T.N., Otto, S., Pham, T., Rimorin, C., Ruiz, A., Sanchez, R., Sawyer, L., Shapovalova, N., Shepard, N., Slaughterbeck, C., Sulc, J., Tieu, M., Torkelson, A., Tung, H., Valera Cuevas, N., Vance, S., Wadhwani, K., Ward, K., Levi, B., Farrell, C., Young, R., Staats, B., Wang, M.Q.M., Thompson, C.L., Mufti, S., Pagan, C.M., Kruse, L., Dee, N., Sunkin, S.M., Esposito, L., Hawrylycz, M.J., Waters, J., Ng, L., Smith, K., Tasic, B., Zhuang, X., Zeng, H.: A high-resolution transcriptomic and spatial atlas of cell types in the whole mouse brain. Nature 624(7991), 317–332 (Dec 2023)

39. Zhang, Z., Sun, H., Mariappan, R., Chen, X., Chen, X., Jain, M.S., Efremova, M., Teichmann, S.A., Rajan, V., Zhang, X.: scMoMaT jointly performs single cell mosaic integration and multi-modal bio-marker detection. Nat. Commun. 14(1), 384 (Jan 2023)

